# Reannotation of eight Drosophila genomes

**DOI:** 10.1101/350363

**Authors:** Haiwang Yang, Maria Jaime, Maxi Polihronakis, Kelvin Kanegawa, Therese Markow, Kenneth Kaneshiro, Brian Oliver

## Abstract

The sequenced genomes in the Drosophila phylogeny is a central resource for comparative work supporting the understanding of the *Drosophila melanogaster* non-mammalian model system. These have also facilitated studying the selected and random differences that distinguish the thousands of extant species of Drosophila. However, full utility has been hampered by uneven genome annotation. We have generated a large expression profile dataset for nine species of Drosophila and trained a transcriptome assembly approach on *Drosophila melanogaster* to develop a pipeline that best matched the extensively curated annotation. We then applied this to the other species to add tens of thousands of new gene models per species. We also developed new orthologs to facilitate cross-species comparisons. We validated the new annotation of the distantly related *Drosophila grimshawi* with an extensive collection of newly sequenced cDNAs. This reannoation will facilitate understanding both the core commonalities and the species differences in this important group of model organisms.

## Background

*Drosophila melanogaster* is a genetic and genomic workhorse that has led to the understanding of chromosome theory of inheritance, the nature of mutations, pattern formation in development, innate immunity, circadian rhythms, and a host of other discoveries in the last century (Bilder and Irvine 2017; Callaway and Ledford 2017). There is a core set of 12 sequenced and assembled genomes in the Drosophila genus (Adams et al. 2000; Richards et al. 2005; Drosophila 12 Genomes et al. 2007; Hoskins et al. 2015). This is an important resource for studying diverse evolutionary biology problems, such as sex chromosome evolution (Charlesworth and Charlesworth 2005), *de novo* gene formation (Lu et al. 2008), and duplication and divergence (Vieira et al. 2007). Using other Drosophila species for comparative genomics can also help identify the conserved genomic elements in *Drosophila melanogaster*, in cases where frequent random occurrence obscures identification of DNA elements (Chen et al. 2014). For example, comparative genomics was a valuable tool for studying Doublesex DNA binding site function, as the short degenerate sequences bound by Doublesex appear by chance at a high rate (Clough et al. 2014). Comparative genomics is also essential for determining the probable function of transcribed elements. For example, short open reading frames (ORFs) in “non-coding” RNAs are not commonly annotated because they occur often in random sequence. But if a short ORF appears in a phylogeny, then those “non-coding” RNAs are likely to encode short biologically active peptides (Tautz 2009).

The utility of the sequenced and annotated Drosophila genomes is clear, but there is room for improvement. The current annotations of *non-melanogaster* members of the genus are uneven and inferior to the heavily and actively curated *D. melanogaster* annotation (Gramates et al. 2017). For example, while there have been six versions of the *D. melanogaster* genome and upwards of 75 annotations (Hoskins et al. 2015), the majority of the other species have a single assembly and one or two annotation versions (Richards et al. 2005; Drosophila 12 Genomes et al. 2007; Hu et al. 2013). Much of genome annotation depends on identification of conserved long open reading frames, but expression data presents a direct way to determine what portions of the genome are actively transcribed and should be annotated.

Dissected adult tissues are a good source for mRNAs to support genome annotation (Chen et al. 2014). While most tissues are present in both sexes, there are some sex-specific organs that show unique expression profiles (Arbeitman et al. 2002; Parisi et al. 2004; Graveley et al. 2011; Brown et al. 2014). For example, there are approximately 8000 genes preferentially expressed in the testis and male reproductive tract and approximately 5000 genes preferentially expressed in ovary and female reproductive tract. Additionally, because female transcripts maternally deposited in eggs are used during embryogenesis, many developmentally important transcripts are detected in adult female ovary samples. Overall, using dissected tissues from adults increases coverage compared to whole samples, due to the fact that genes rarely expressed in a whole organism often show enriched expression in a given tissue (Chintapalli et al. 2007). Greater than 85% of annotated genes are expected to be covered in such experiments (Chintapalli et al. 2007; Daines et al. 2011). In this work, we have used Poly-A^+^ RNA-seq expression profiling of 584 samples from adults of *D. melanogaster (Dmel), D. yakuba (Dyak), D. ananassae (Dana), D. pseudoobscura (Dpse), D. persimilis (Dper), D. willistoni (Dwil), D. mojavensis (Dmoj), D. virilis (Dvir)*, and *D. grimshawi (Dgri)* to support reannotation of the corresponding genomes.

Performing *de novo* annotation based on gene expression is complicated by RNA coverage gaps that result in discontinuity within a single transcription unit, overlapping genes, and false splice junction calls due to gap generation that maximizes read alignment (Robertson et al. 2010; Sturgill et al. 2013). As a result, reference annotations and *de novo* transcript assemblies can differ radically (Garber et al. 2011; Haas et al. 2013). Some of these difficulties can be overcome, for example by using methods that capture strandedness (Grabherr et al. 2011). Additionally, tuning the transcript parameters can improve the quality of the transcriptome (Vijay et al. 2013). These tools are often run using default parameters or based on some simple assumptions and tests. In this work, we decided to systematically test parameters and train support vector machines on *D. melanogaster*, and then lift over these settings for automated annotation of the remaining species. This resulted in dramatic improvements in the mapping of RNA-seq reads to a greatly expanded set of genes and isoforms in these species.

## Results and Discussion

### RNA-seq

*Dmel, Dyak, Dana, Dpse, Dper, Dwil, Dmoj, Dvir*, and *Dgri* is a wide range of species separated by an estimated 40 million years of evolution (Leung et al. 2015), with fully saturated neutral substitutions at the widest separations (Figure 1A; after (Chen et al. 2014)). To evaluate the annotations of these nine members of the genus, we performed stranded Poly-A^+^ RNA-seq experiments on quadruplicate biological samples derived from sexed whole flies and tissues (covering up to eight adult tissue types for each sex: whole organism, gonad, reproductive tract, terminalia, thorax, viscera, head, and obdomen) for a total of 584 samples and ~5 billion RNA-seq reads (available at the Gene Expression Omnibus, GEO (Edgar et al. 2002), under accession GSE99574 and a subset of accession GSE80124, see Methods). We targeted the *Dyak, Dana, Dpse, Dper, Dwil, Dmoj, Dvir*, and *Dgri* genomes for reannoation. We also included two *Dmel* strains; *w*^*1118*^ and *Oregon-R (OreR)* to facilitate training the annotation pipeline.

**Fig. 1.**
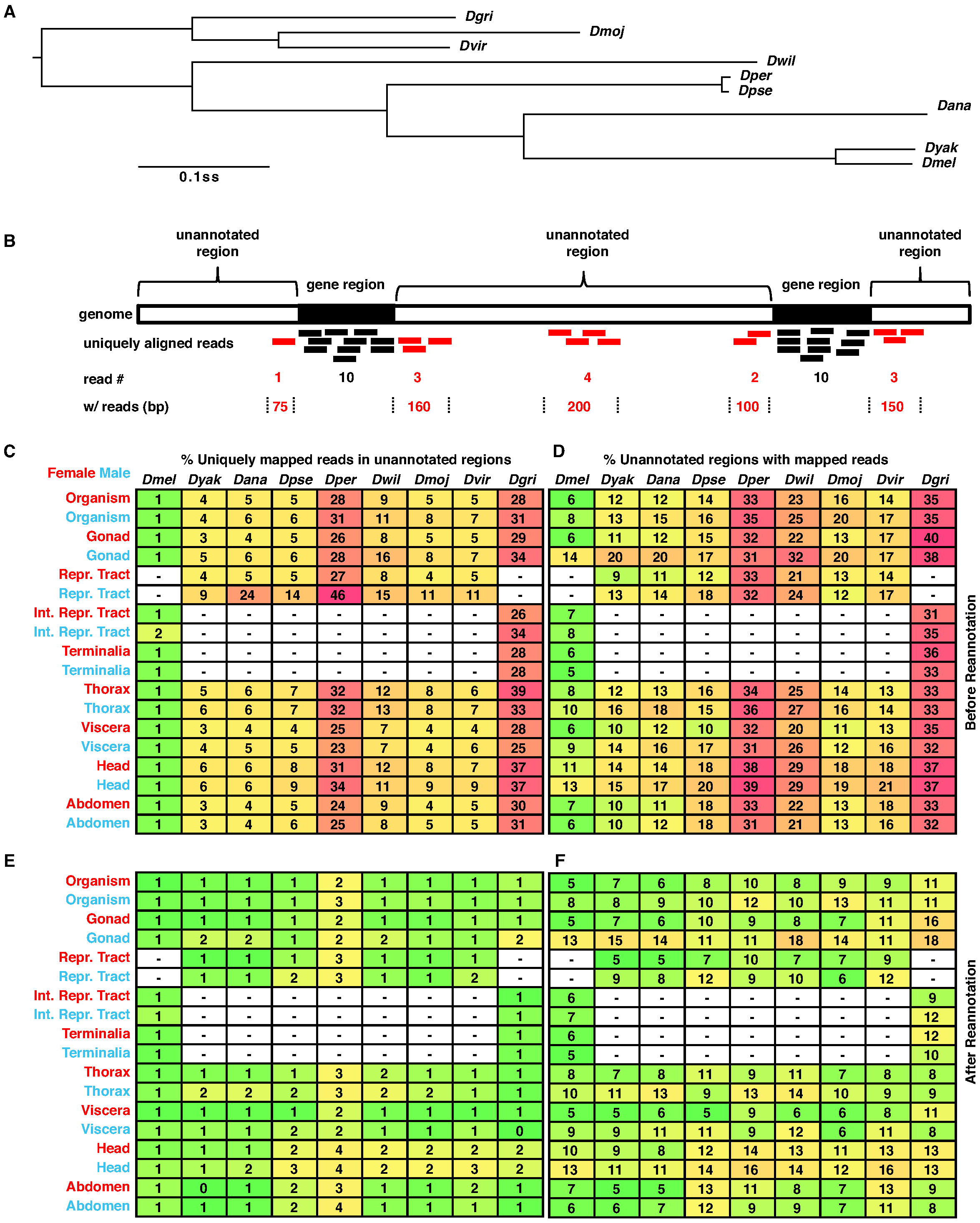
Evaluation of reannotation for non-melanogaster Drosophila species. (A) Bayesian phylogenetic tree of nine Drosophila species. Nodes are supported by 100% posterior probabilities, and phylogenetic distance is shown as substitutions per site (ss). After (Chen et al. 2014). Standard three letter abbreviations are used (see text). (B) Cartoon of measurements of reads mapped to unannotated regions. Gene regions (fill), unannotated regions (open), RNA-seq reads (bars) mapped to gene regions (black) or unannotated regions (red) are shown. Numbers of reads (#) and regions (in bps, flanked by dotted lines) with reads are also shown. Percentage of reads (C, E) and regions with mapped reads (D, F) in unannotation regions before (C, D) and after (E, F) reannotation. Green, yellow, and red denotes low, median, and high percentage, which are also indicated in each cell. Expression profile tissues are shown (left) for female (red) and male (blue) samples. Reproductive (Repr.) tract includes the internal ducts and glands and the terminalia. Internal Reproductive (Int. Repr.) tract does not include the Terminalia. Missing tissues are shown (dash and open fill).

### Annotation evaluation and optimized reannotation

We first used RNA-seq data to evaluate the existing annotations. If an annotation was complete, the vast majority of our RNA-seq reads would map to annotated transcripts. On the other hand, if an annotation was poor, we would observe more RNA-seq reads mapping to unannotated regions and there might be extensive unannotated regions with mapped reads (Figure 1B). To determined how many reads aligning to the genomes mapped to annotated genes and transcripts, and how well those reads covered the existing models, we calculated the ratio of reads uniquely mapped to unannotated regions relative to the ones uniquely mapped to the whole genome by tissue type and species (Figure 1C). This metric was sensitive to read abundance from highly expressed unannotated genes. Therefore, in addition to the number of RNA-seq reads, we used a related metric describing the number of bases covered by at least one RNA-seq read outside of the annotations (Figure 1D).

We observed a wide range of annotation qualities using these simple metrics. For example, *Dmel* had only 1% of reads uniquely mapped to unannotated regions for all tissues except male reproductive tract, which had 2% mapping to unannotated regions, strongly suggesting that the *Dmel* genome annotation for highly expressed genes is nearly complete (Figure 1C). However, we still observed read alignment at up to 14% of unannotated *Dmel* regions (Figure 1D), suggesting that additional annotation is required, especially for fully capturing the transcriptomes of the testis, head and thorax. In contrast, we observed that 3-46% of reads uniquely mapped to unannotated regions in *non-melanogaster* species (Figure 1C). Similarly, we observed that 9-40% of unannotated regions had mapped read coverage (Figure 1D). *Dper* and *Dgri* had the poorest annotations. To determine if certain tissues might be especially valuable in completing the transcriptome, we also examined the reads mapping ratios by tissue. As we observed for *Dmel*, we found that RNAs from ovary, reproductive tract, thorax, viscera, and abdomen had the best mapping, while RNAs from testis and head of either sex had the poorest mapping to the annotations. In conclusion, our results suggested that all eight of the non-melanogaster annotations need major improvements to approach the quality of the *Dmel* annotation.

Given the clear superiority of the *Dmel* annotations from the read mapping, we decided to systematically develop *de novo* transcriptome annotations for the genus that would approximate the *Dmel* annotation quality. In order to determine the best method for generating these new annotations, we generated tens of thousands of *de novo* annotations of *Dmel*, where we systematically and iteratively honed-in on optimal transcript assembly algorithm parameters, and used support vector machines to develop filtering criteria, such that we most closely matched the *Dmel* standard. We then used these settings and filters to generate new annotations for the remaining species (see Methods). We were gratified to find that in non-melanogaster species, the reads mapping to unannotated regions decreased five-fold after the annotation update (Figure 1E,F; Wilcoxon rank test, *p* < 2.2×10^−16^). As expected based on our use of *Dmel* as a “gold standard”, there was no significant improvement in reads mapping exclusively to the annotation in *Dmel.* This harmonized annotation will provide an improved basis for comparative genomics studies.

### Summary of new gene models

Because the original annotation pipelines for the non-*melanogaster* members focused on conserved longest ORFs at each locus (Drosophila 12 Genomes et al. 2007), we anticipated that much of the improvement to the annotation would come from extending the annotation of untranslated regions (UTRs) and new isoforms due to alternative promoters, termination, and alternative splicing, as well as noncoding or minimally coding RNAs (ncRNAs). Indeed, we found that ~8K new gene models per species overlapped with and extended the older annotations (Figure 2A). For example, the *Dwil* gene *GK27243*, which is expressed in the testis, had the same splice junctions in the old and new annotation *(YOgnWI09161)*, but had longer 5’-and 3’-ends in the updated annotation (Figure 2B). We also observed an increase of 10-20K isoforms in the new annotation compared to the old one (Figure 2C). For example, the *Dper doublesex (dsx)* locus *(GL23549)* had a single annotated isoform (Figure 2D), whereas in *Dmel* the *dsx* locus encodes sex-specific transcription factors from sex-specifically spliced pre-mRNAs (Burtis and Baker 1989). Our new annotation *(YOgnPE00925)* captures sex-specific isoforms of *Dper dsx* and includes a new upstream promoter. We also observed 0.7-1.3K instances per species where gene models were merged (Figure 2E). In at least some cases, this was strongly supported by expression data. For example in the case of the *YognWI03804* locus, the last two exons of *Dwil GK26840* are clearly joined by junction reads to the single exon of *GK20038* locus forming an updated gene model (Figure 2F). However, we did observe ~700 instances of merging in the well-annotated *Dmel* genome, which seems excessive. We generated 1-2K completely novel annotations per species (Figure 2G,H). These included ncRNAs (~24% of novel annotations), such as the *Dyak* non-coding homolog *(YOgnYA12879)* of *rna on X 1 (roX1). Dmel roX1* is a component of the male-specific X-chromosome dosage compensation complex (Kuroda et al. 2016), and like the *Dmel* ortholog, the *Dyak roX1* locus is expressed in males, but not females. Loci producing ncRNAs tend to diverge rapidly, but both the *Dmel* and *Dyak roX1* loci are flanked by the *yin* and *echinus* orthologs. The combination of sequence, expression pattern, and synteny strengthen the conclusion that these *roX1* genes descended from a common ancestral gene.

**Fig. 2.**
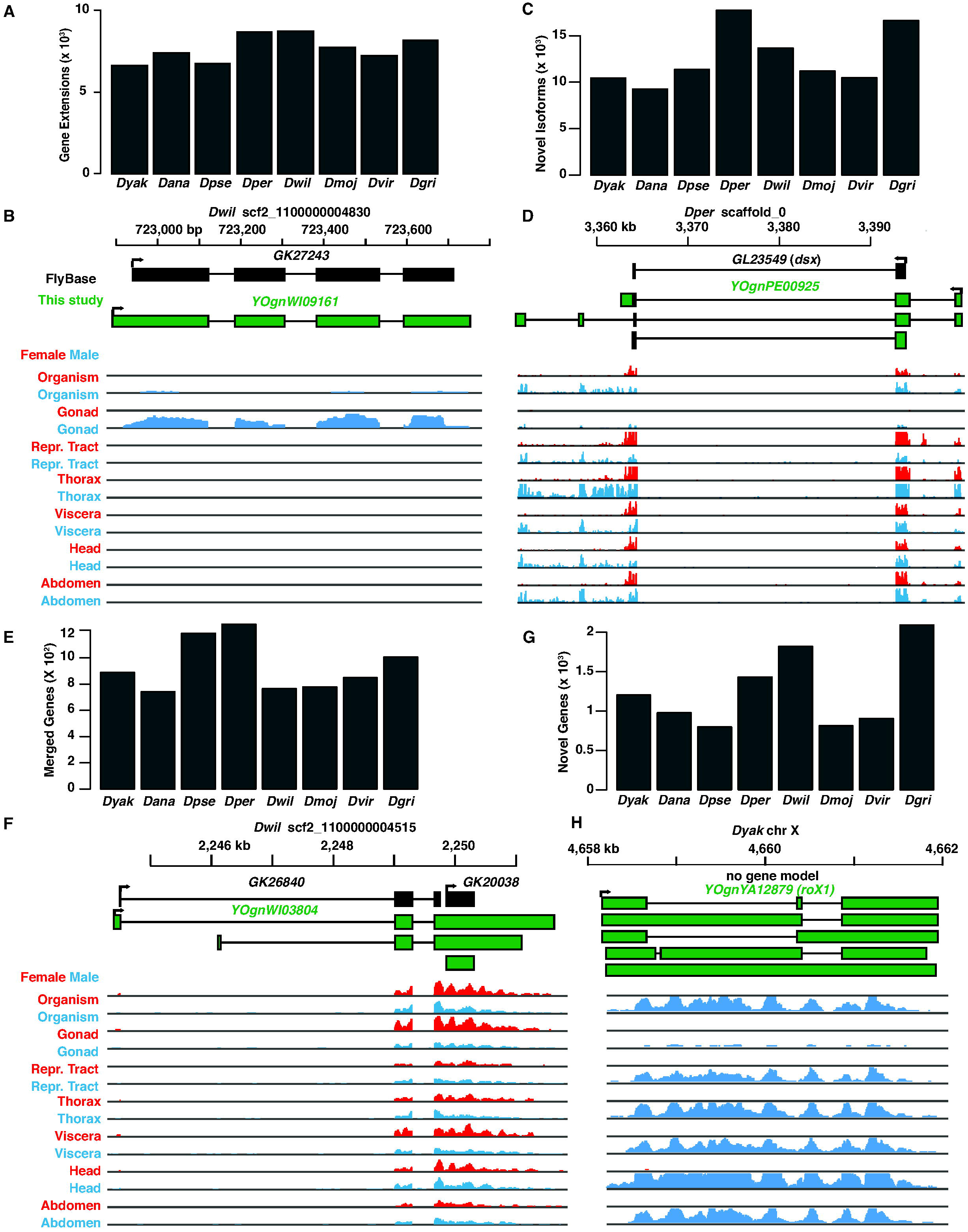
Summary and reannotation. (A) Number of new gene annotations with extended coverage and (B) one example. (C) Number of novel isoforms in the reannotation and (D) one example. (E) Number of merged genes in the reannotation and (F) one example. (G) Number of novel genes in the reannotation, and (H) one example. (A-H) Species, scaffold IDs, and locus names are shown. FlyBase gene models (black) and new models (green) are shown. Orientation of transcripts are shown (arrow at 5’-end). Expression level tracks (arbitrary FPKM scale) for indicated tissues/sexes are shown.

To identify other novel orthologs like *roX1*, we analyzed the synteny, sex-and tissue-biased expression patterns, and gene structures of previously identified orthologs of *Dmel* genes and developed a support vector machine to generate a list of candidate orthologs, which were then compared at the sequence similarity level (see methods). We called 500-1000 new orthologs per species (Figure 3A). For example, we found orthologs of the non-coding *Dmel CR42860* gene in four of the eight other species *(YOgnYA06038, YOgnAN10714, YOgnWI07915*, and *YOgnVI13637* in *Dyak, Dana, Dwil*, and *Dvir* respectively; Figure 3B). In each case, the gene is most strongly expressed in the thorax. Interestingly, in *Dwil, YOgnWI07915* also showed female-biased expression (*p*_*ajd*_ = 5.4×10^−10^, DESeq2), highlighting the fact that we can observe changes in sex-and/or tissue-biased expression in the phylogeny. Extending the Drosophila orthology to include ncRNAs should allow for the exploration of conserved and divergent functions of this understudied aspect of comparative genomics.

**Fig. 3.**
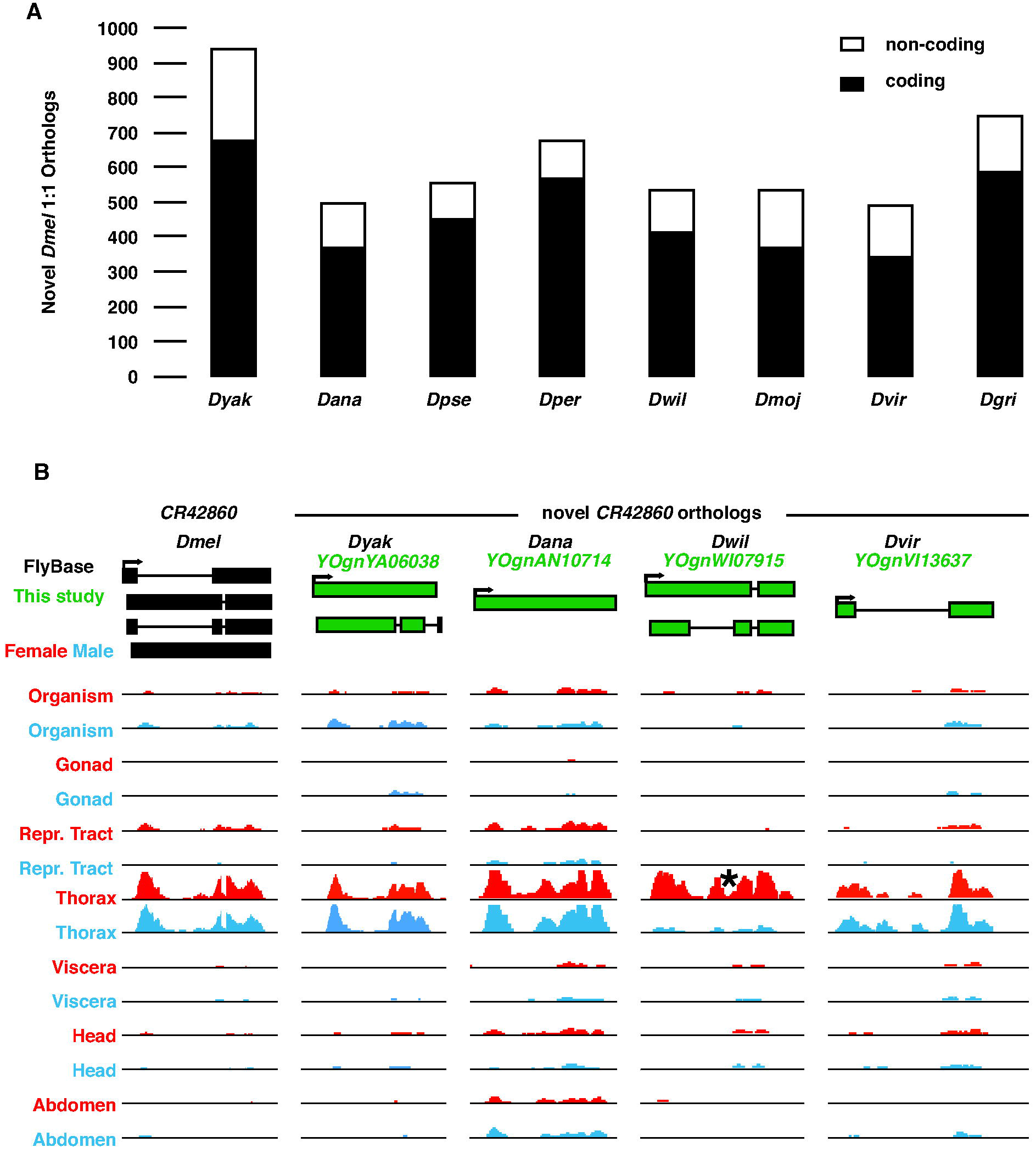
Summary and novel 1:1 orthologs. (A) Number of novel *Dmel* 1:1 orthologs in each of the eight non-melanogaster species. The coding (filled) and non-coding (open) status of genes in *Dmel* are. (B) An example of novel 1:1 orthologs of *Dmel CR42860* identified in *Dyak, Dana, Dwil*, and *Dvir. CR42860* is nested in the intron of sls in all five species. Significant (*p*_*adj*_ = 5.4×10^−10^) female-biased expression of *Dwil* ortholog is shown (asterisk). See Figure 2 for abbreviations and color coding.

### cDNA validation

Since we used the illumina RNA-seq data to build the new annotations, we needed an independent transcriptome dataset for validation. The *Dgri* annotation was greatly changed in our work and has been unexplored at the RNA-seq level. We therefore chose to validate a subset of the *Dgri* annotation. Longer reads have a distinct advantage for this validation, as they can capture the alternative splice forms without reliance on short junction reads. Therefore, we conducted duplicate PolyA+ PacBio Iso-seq for *Dgri* sexed adults. To better sample the transcriptome, we fractionated each RNA preparation into three size categories (see Methods). We generated 282,616 high quality, full length, non-chimeric, cDNAs and compared overlap with the annotation. This sampling experiment covered 27-29% of the FlyBase and new annotations (Figure 4A).

**Fig. 4.**
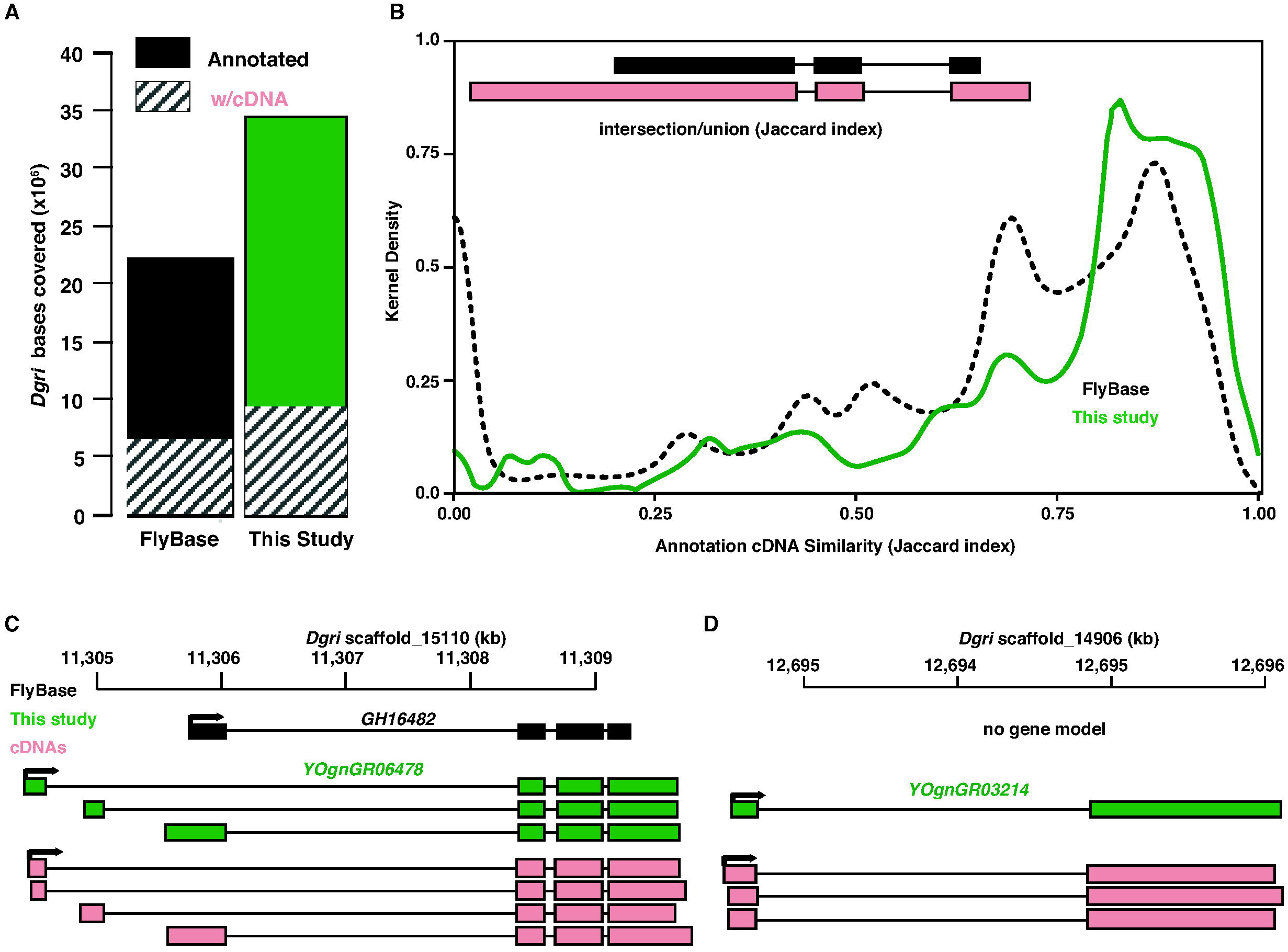
PacBio cDNA validation of *D. grimshawi* annotation. (A) Bases in the genome covered by annotations (whole bars) and validated by PacBio Iso-seq cDNA sequencing (stripes) in *Dgri.* (B) Similarity between annotated gene models and cDNAs before (dotted black line) and after reannotation (solid green line) as measured by Jaccard index (intersection / union), as illustrated (inset). Jaccard index distributions are shown in kernel density. (C) An example of cDNA (SRR6840922) aligning to a previous and new gene model, where the new gene model have extended coverage and a greater Jaccard index. (D) An example of cDNAs (SRR6840922) aligning with a novel gene model and a jaccard score of “0” with the previous annotation. See Figure 2 for abbreviations and color coding.

To systematically analyze the relationship between the cDNA sequences and the annotations, we measured the intersection/union (Jaccard index) for base coverage at the transcript isoform level (Figure 4B). The most dramatic difference in the Jaccard index was at zero, where there is no overlap between cDNAs and annotations. We observed a dramatic decrease in the fully non-overlapping cDNAs when we used the new annotation. The distribution of Jaccard index scores shows a shift towards fully overlapping, with 56% of cDNA/new annotations showing 75-100% similarity. For example, four cDNAs mapping to the *Dgri GH16482* locus (Jaccard index >0.94) validated the three isoforms of *YOgnGR06478*, with three distinct promoters, each of which has an extended 3’-UTR relative to the FlyBase annotation (Figure 4C). Similarly, we found support (Jaccard index >0.95) for the previously unannotated locus *YOgnGR03214* (Figure 4D).

## Conclusions

Previous work has used expression data in the Drosophila genus to validate the gene models in *Dmel* (Drosophila 12 Genomes et al. 2007; Chen et al. 2014), but there has been less systematic effort to use the knowledge from *Dmel* to inform and annotate the rest of the genus. To maximize the value of the sequenced Drosophila genomes, we have generated an extensive expression profile in order to assemble transcript models and update the annotations. In all cases this resulted in an extensive set of predicted transcripts. To leverage the decades of dedicated annotation that has been performed on *Dmel* (Adams et al. 2000; Celniker 2000; Lewis et al. 2002; Misra et al. 2002; Celniker and Rubin 2003; Drysdale 2003; Ashburner and Bergman 2005; Brown and Celniker 2015), we generated thousands of *de novo Dmel* annotations, to determine optimal parameters and filters that resulted in the best match to the existing *Dmel* annotation. We then applied this optimized pipeline to the rest of the genomes. This was largely successful, as we not only generated tens of thousands of new isoform models in each species, but we were also able to validate these models in *Dgri* with an independent set of full length cDNAs. We suggest that when there is a high-quality annotation of a given species, this methodology could be used to tune annotation pipelines for related species. The fact that this worked well for a species, *Dgri*, that is separated from *Dmel* by about 40 million years (Leung et al. 2015), suggests that targeting a few genomes in a lineage for full curation, can then result in a high-quality annotation for scores of related species.

Given how often Drosophila are used in evolutionary studies on gene birth, death, duplication, and divergence, having improved annotations will facilitate a large number of studies. Similarly, workers interested in understanding *Dmel* genes should find these annotations useful for determining if a given feature is evolutionarily conserved. For example, this should be particularly true for identifying ncRNAs and protein binding sites in UTRs, as those features were lacking for the majority of non-*Dmel* genes. To make these annotations useful we have posted gene models and data in a number of formats. These annotations will be relatively easy to update, without new experiments, if the underlying assemblies are updated, as we saved all the unmapped reads at the sequence read archive (Leinonen et al. 2011).

## Methods

See the extensive reagent and resources table for strains, media, reagents and suppliers, software, database submissions, and other identifiers (Additional File 1).

### Flies

Flies were grown at the NIH *(Dmel)*, the UCSD Species Stock Center (*Dyak, Dana, Dpse, Dper, Dwil, Dmoj*, and *Dvir*), or the Hawaiian Drosophila Research Stock Center (*Dgri*). Growth conditions are given in GEO GSE99574 and GSE80124. For each sex and species, we dissected seven to eight adult tissues or body parts in PBS and transferred the samples immediately to RNAlater.

### RNA-seq

Illumina sequencing details are given in GEO GSE99574 and GSE80124. Briefly, we isolated RNA using the RNeasy 96 kit. We added ERCC spike-ins (Jiang et al. 2011; Zook et al. 2012) for quality control purposes. We conducted single-end stranded 76 bp polyA+ RNA-seq experiment for all samples using the TruSeq kit and protocol. We quantified nucleic acids with Quant-iT RiboGreen or PicoGreen kits. We multiplexed using both index adaptors and mixing RNA from distantly related species and sequenced on the HiSeq2000 Sequencing System. De-multiplexed reads were produced by Illumina CASAVA (v1.8.2) as fastq files. We mapped the reads of mixed species libraries with HiSAT2 (v2.0.5; --dta and-max-intronlen = 300,000) and used SAMtools (v0.1.19) (Li et al. 2009)to sort the HiSAT2-generated bam files by read name. We used a python script demultiplexer (v1.0) to scan the bam file to collect the reads specific to one species or ERCC spike-ins (Jiang et al. 2011; Zook et al. 2012). We converted from bam to fastq format using BEDTools (v2.25.0; bamtofastq) (Quinlan and Hall 2010). We used the most current annotations of the species (FlyBase release 2017_03), except for *Dgri* because NCBI and FlyBase used our pre-publication RNA-seq data to improve the annotation of *Dgri* using gnomon (Kapustin et al. 2008). We used the the last version of *Dgri* annotation, prior to RNA-seq data inclusion (Flybase release 2016_05).

PacBio cDNAs details are found at NCBI SRA (SRP135764). Briefly, we constructed *Dgri* cDNA libraries following the Isoform Sequencing (Iso-Seq) protocol using the Clontech SMARTer cDNA Synthesis Kit and the SageELF size-selection system. We used 500 ng total RNA per reaction for the polyA+ enrichment and first strand synthesis, and conducted the first round of PCR amplification (95 °C for 2 min; 14 cycles of 98 °C for 20 sec, 65 °C for 15 sec, 72 °C for 4 min; 72 °C for 5 min) to generate double strand cDNA for size selection. We used three fraction ranges (SageELF index 10-12 or 1-2 kb, 8-9 or 2-3 kb, and 5-7 or 3-5 kb) of double strand cDNA for the second round of PCR amplification (95 °C for 2 min; N cycles of 98 °C for 20 sec, 65 °C for 15 sec, 72 °C for X min; 72 °C for 5 min). We repaired DNA ends and performed blunt-end ligation. We quantified SMRTbell libraries by Qubit Fluorometric Quantitation and qualified by Bioanalyzer beforesequencing on the PacBio RS II using DNA Sequencing Reagent kit 4.0 v2 with a run time of 240 minutes. We used 6th generation polymerase and 4th generation chemistry (P6-C4). Circular consensus (ccs2) reads (-- maxLength=40000 --minPasses=1) were generated using PacBio pitchfork (v0.0.2) after conversion of bax.h5 files to bam using bax2bam. The final full length non-chimeric Iso-seq reads were concatenated from three fractions and available in SRA (SRP135764).

### Annotation optimization

We developed a pipeline to match the existing *Dmel* annotation with *de novo* RNA-seq data (Figure 5A). We mapped reads with HiSAT2 (v2.0.5; --dta and-max-intronlen = 300,000) (Kim et al. 2015). We then used StringTie (v1.3.3) (Pertea et al. 2015) to generate *de novo* annotation using the bam alignments from HiSAT2. We set minimum transcript length according to the shortest gene in *Dmel* (i.e., 30 bp), and we set the strandedness library to “-rf”. We optimized “-c” (minimum reads per bp coverage to consider for transcript assembly), “g” (minimum gap between read mappings triggering a new bundle), “-f” (minimum isoform fraction), “-j” (minimum junction coverage), “-a” (minimum anchor length for junctions), and “-M” (maximum fraction of bundle allowed to be covered by multi-hit reads). We used a union set of reads from replicated *w*^1118^ and *OreR* females and males to optimize StringTie parameters. To test which combination of parameters generated the *de novo* annotation closest to FlyBase, we used the Jaccard index (BEDTools v2.25.0) of unique exons to measure similarity. In the first round of testing (Figure 5B), we tested all combinations of “c” (1, 3, 5, 7, 9), “g” (10, 30, 50, 70, 90), “f” (0.01, 0.03, 0.05, 0.07, 0.09), “j” (1, 2, 3, 5, 7, 9), “a” (5, 10, 15, 20, 25), and “M” (0.1, 0.3, 0.5, 0.7, 0.9). Among the 15,625 tests, the parameters with the highest Jaccard index were “c” = 3, “g” = 50, “f” = 0.01, “j” = 3, “a” = 10, and “M” = 0.9. In the second round of testing, we further picked all combinations of the points next to the previous optimal parameters with smaller intervals-“c” (1, 1.5, 2, 2.5, 3, 3.5, 4, 4.5, 5), “g” (30, 40, 50, 60, 70), “f” (0.005, 0.01, 0.015, 0.02, 0.025, 0.03), “j” (1, 2, 3, 4, 5), “a” (5, 6, 7, 8, 9, 10, 11, 12, 13, 14, 15), and “M” (0.7, 0.75, 0.8, 0.85, 0.9, 0.95). Among the 89,100 tests, the parameters with the highest Jaccard index were “c” = 1.5, “g” = 50, “f” = 0.015, “j” = 1, “a” = 14, and “M” = 0.95. In the third round of test, we further picked all combinations of the points next to the previous optimal parameters with smaller intervals-“c” (1, 1.1, 1.2, 1.3, 1.4, 1.5, 1.6, 1.7, 1.8, 1.9, 2), “g” (40, 41, 42, 43, 44, 45, 46, 47, 48, 49, 50, 51, 52, 53, 54, 55, 56, 57, 58, 59, 60), “f” (0.01, 0.011, 0.012, 0.013, 0.014, 0.015, 0.016, 0.017, 0.018, 0.019, 0.02), “j” (1, 2), “a” (13, 14, 15), and “M” (0.9, 0.91, 0.92, 0.93, 0.94, 0.95, 0.96, 0.97, 0.98, 0.99). Among the 152,460 tests, the parameters with the highest Jaccard index were “c” = 1.5, “g” = 51, “f” = 0.016, “j” = 2, “a” = 15, and “M” = 0.95. We applied the optimized parameters for all Drosophila species to generate sample-level *de novo* annotations. We then merged these sample-level annotation to species-level annotation for each species by StringTie (--merge). We optimized three parameters-“-F” (minimum input transcript FPKM), “-T” (minimum input transcript TPM), and “-g” (gap between transcripts to merge together) using the same optimization pipeline (Figure 5C). The optimal parameter combination was F=0, T=10, and g=0. We set minimum input transcript coverage (“-c”) and minimum isoform fraction (“-f”) as 1.5 and as 0.016 respectively to be consistent with previous StringTie settings. The effect of optimization on output is illustrated in Figure 5D. We applied the optimized parameters for all Drosophila species to generate draft species-level *de novo* annotations.

**Fig. 5.**
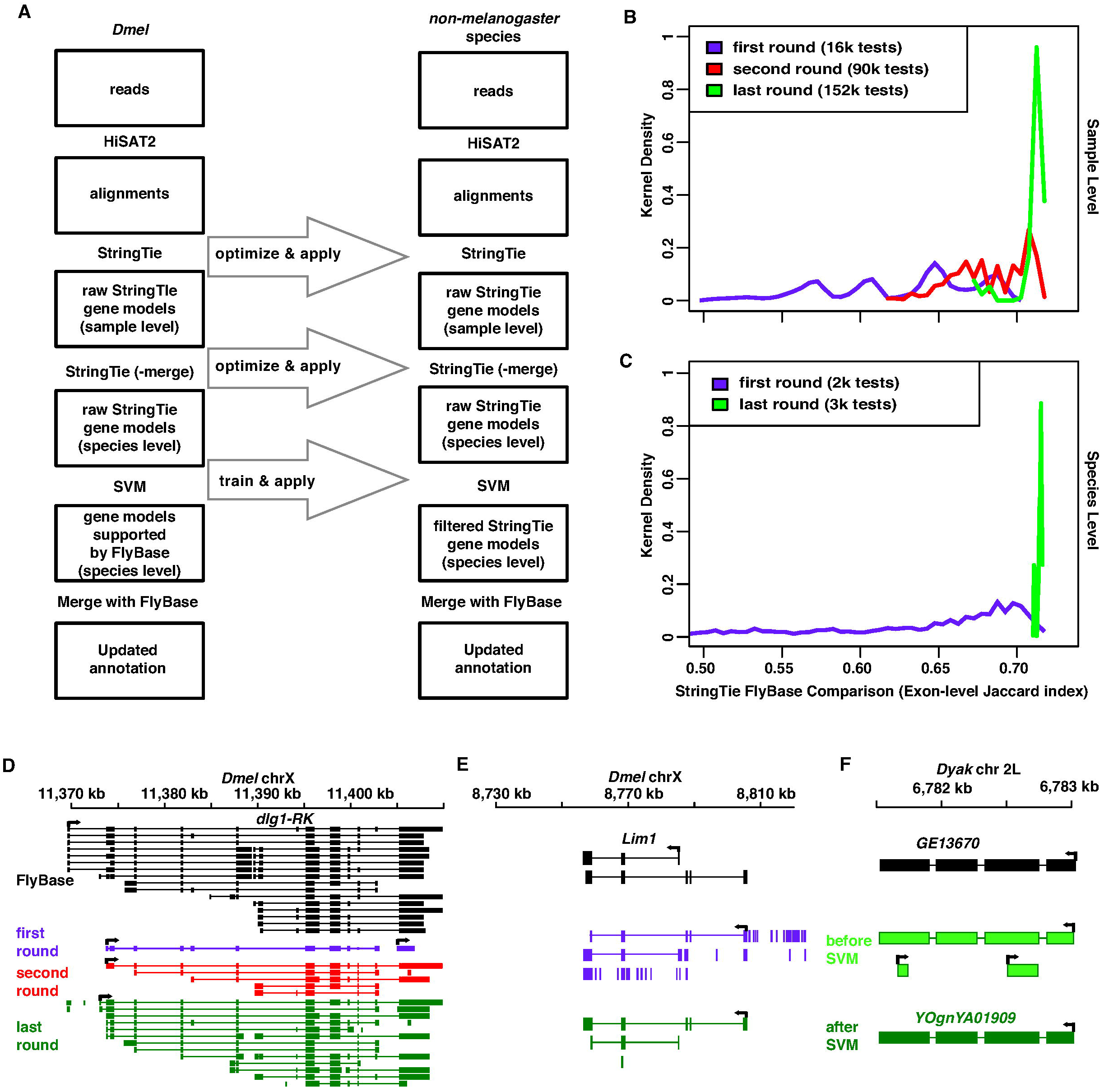
Reannotation pipeline and summaries. (A) Reannotation pipeline. From top to bottom, HiSAT2 was used to map all RNA-seq reads back to appropriate genome. StringTie was used to generate sample-level annotations as well as merging annotations to the species-level. Support Vector Machines (SVM) were used to train the recognition of FlyBase gene models based gene features (e.g. exon length, intron length, and isoforms. See Methods). In each species, the new annotation was eventually merged with the corresponding FlyBase annotation by gffcompare to create the new annotation. The same optimized parameters in *Dmel* (left column) were applied in each of the non-melanogaster species (right column). (B) In *Dmel*, we progressively converged sample level annotations on the FlyBase annotation in three rounds of parameter optimization in StringTie. (C) Again in *Dmel*, we permuted and tested species level annotations to maximize similarity to the FlyBase annotation during parameter optimization in StringTie (-merge). (B,C) The X-axis is the exon-level Jaccard index between StringTie and FlyBase annotation. The Y-axis is the distribution of Jaccard index scores in kernel density. First (purple), second (red), and last (green) rounds of optimization are shown. (D) An example of gene model improvements when generating sample level annotations. (E) An example of gene model improvements when generating species level annotations. Note that most gene models with short, single and intron-less transcripts were removed in the last round of parameter optimization. (F) An example of applying SVM in a non-melanogaster species. FlyBase gene model was shown in black, and Gene models before (light green) and after SVM (dark green) are shown. See Figure 2 for abbreviations and additional color coding information.

To identify which StringTie predicted gene models were already annotated in FlyBase, we used Jaccard index to calculate similarity of gene structure between StringTie gene models and FlyBase. We used a cutoff (Jaccard index > 0.6), to obtain ~10k genes identified by both StringTie and FlyBase, and we deemed these genes as correctly predicted *Dmel* gene models of StringTie. However, we still observed an abundance of short single exon genes called by StringTie but not FlyBase (Figure 5E). To remove more of the StringTie unique calls, we used sequential and expressional features of genes to train a Support Vector Machine (SVM) (sklearn package v0.19.1 of Python v3.4.5) to recognize all the StringTie-predicted gene models with Jaccard index > 0.6 with existing gene models in FlyBase. The features included isoform number, exon size, exon GC%, intron size, intron GC%, intron number (GT-AG intron type and other intron type respectively), and median expression (DESeq2 normalized read counts) in 14 sexed tissues (except terminalia). For the SVM parameters, we tested different kernels (i.e., rbf, sigmoid), penalty parameter C (0.001, 0.01, 0.1, 1, 10, 100, 1000), and kernel coefficient gamma (0.001, 0.01, 0.1, 1, 10, 100, 1000) values. The Receiver operating characteristic (ROC) analyses indicated that the Area Under Curve (AUC) was the largest (0.97) at kernel = rbf and C = 10 and gamma = 0.1. We also tested penalty parameter C (0.001, 0.01, 0.1, 1, 10, 100, 1000) under the linear kernel (kernel coefficient gamma is not available for this kernel), the maximum AUC we obtained is 0.95, smaller than the optimal parameters. We applied the same SVM model with the optimized parameters to find qualified gene models in each species (Figure 5F).

To keep all StringTie gene candidates and FlyBase gene models in the updated annotation, we merged all qualified StringTie gene models with FlyBase annotation using gffCompare (v0.9.8) with option-r. In *Dmel*, we merged the StringTie gene candidates that were identified as correct prediction (i.e., Jaccard index >0.6 to FlyBase gene model) to the FlyBase annotation. If a StringTie transcript and a FlyBase transcript share the same structure for all introns on the same strand, we used the union of the gene structure of StringTie and FlyBase transcripts. After this step, the updated annotations were generated for each species. We used a universal ID format in the final updated annotations (e.g. YOgnYA12345). The format is Yang and Oliver (YO), gene name (gn), species; *Dmel* (ME), *Dyak* (YA), *Dana* (AN), *Dpse* (PS), *Dper* (PE), *Dwil* (WI), *Dmoj* (MO), *Dvir* (VI), and *Dgri* (GR), gene (gn) or transcript (tr), and a numerical identifier.

### Updated orthologs

We used gene synteny, tissue-level expression, splicing conservation, and sequential similarity relative to *Dmel* to search for new orthologs (Figure 6A) using random gene pairs as a null model. To determine flanking gene synteny, we first identified the flanking 10 genes on each side of the target gene determined by the best reciprocal blast hits using blastp (Altschul et al. 1990) of all the known protein sequence in FlyBase (Figure 6B). To compare the expressional similarity, we plotted the normalized read counts of 14 tissues (all except terminalia) in the species gene relative to the *Dmel* gene (Figure 6C). When comparing the gene structure similarity of any gene pair between *Dmel* and non-melanogaster, we calculated the ratio of (unique intron number + 1) between species gene and *Dmel* (Figure 6D). We used all known one-to-one orthologs between *Dmel* and non-melanogaster species to train a SVM model to recognize possible orthologs candidates based on the above three features. The optimal parameter combination were rbf kernel, C = 1000, and gamma = 0.001. This procedure created a large number of potential orthologs. To be more conservative, we further filtered these ortholog candidates by using sequential similarity. We aligned the longest transcript of *Dmel* genes with that of non-melanogaster ortholog candidate by ClustalW (v2.1) with default parameters (Thompson et al. 1994). For each alignment, we calculated the ratio of alignable length to total length, and for the alignable region we further calculated the ratio of identical length to alignable length. We used these two features to run SVM model (kernel = rbf, C = 1000 and gamma = 0.01) and selected the predicted orthologs (Figure 6E).

**Fig. 6.**
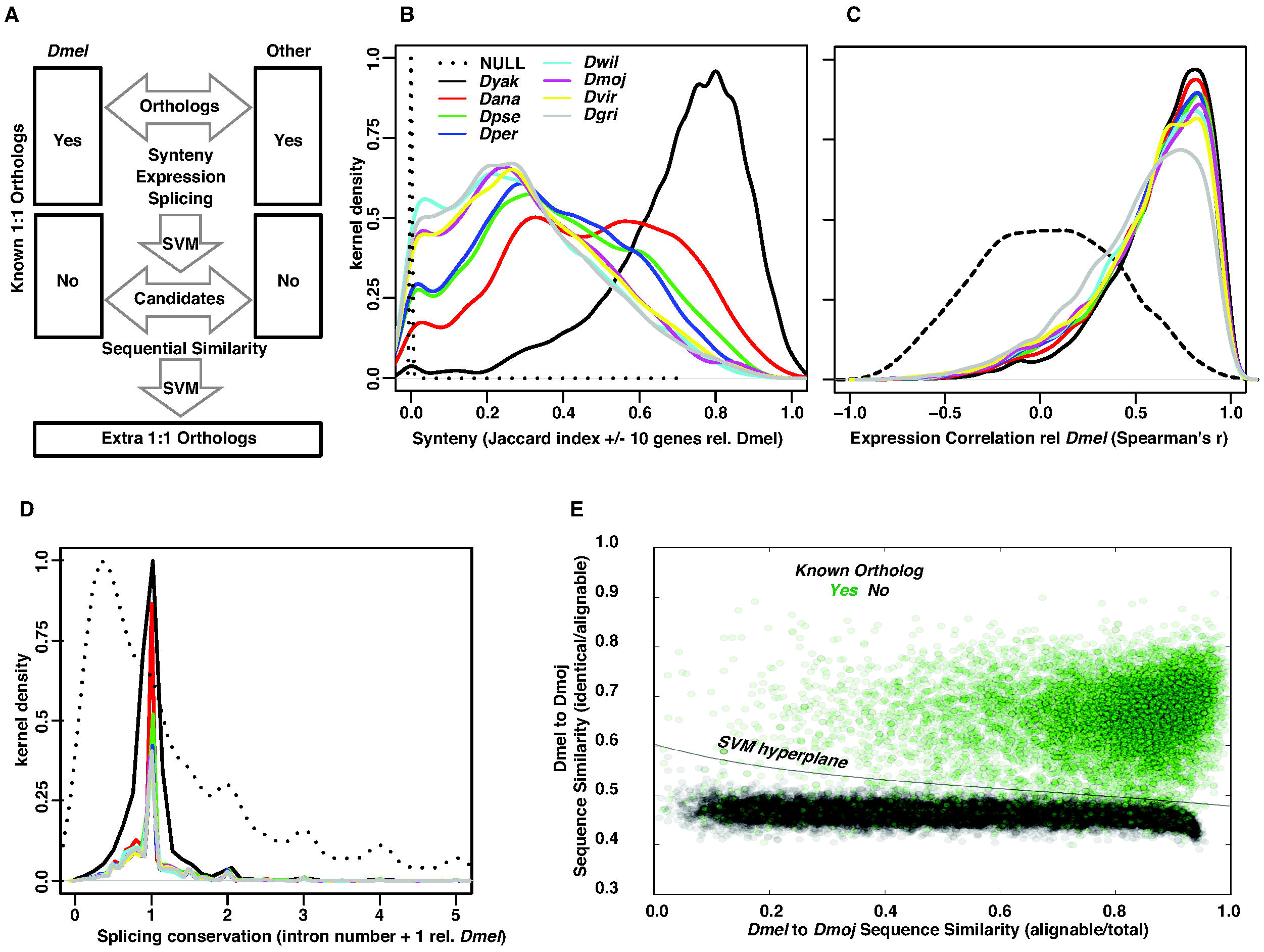
Ortholog identification pipeline and summaries. (A) Pipeline to obtain extra 1:1 orthologs relative to *Dmel.* We used gene synteny, expression correlation among tissues, and exon structure similarity to train SVM models to recognize all known 1:1 orthologs. We used the same SVM model to generate more ortholog candidates among the genes that were not included in the current 1:1 ortholog dataset. Then we used sequence similarity (both alignable/total and identical/alignable) to finalize the extra 1:1 orthologs. (B) Distribution of orthologs in +/-10 gene window surrounding the query gene relative to *Dmel* for each non-melanogaster species (in kernel density). Solid lines are distributions of previously reported 1:1 orthologs, and dotted line (NULL group) is the expected distribution based on random gene pairs (generated by python random package) between *Dmel* and *Dyak* (the other non-melanogaster species all generate identical distributions) in genome (the same for the following panels). (C) Distribution of expression similarity among orthologs (Spearman’s r) relative to *Dmel* for each non-melanogaster species (in kernel density). For each ortholog, normalized read counts of 14 sexed tissues between *Dmel* and each non-melanogaster species were used to calculated correlation. (D) Distribution of intron number relative to *Dmel* for each non-melanogaster species. Intron number plus one was used to avoid infinit value. (E) An example of SVM training of sequence similarity between *Dmel* and *Dmoj.* Known orthologs were shown in green dots, while random gene pairs were shown in black dots. The SVM hyperplane were shown in solid line.

## Supplementary Files

All data resouces and their locations were listed in the Reagent Table (Additional File 1). Briefly, the following is available at GEO under GSE99574 and GSE80124 (see specific GSM#s in Additional File 1): original RNA-seq reads (in fastq format); two demultiplexing versions, one based on HiSAT2 used here, and another generated by STAR; HTSeq raw read counts; gene-level DESeq2 normalized read counts; transcript-level expression from Salmon based on the updated annotation; bigWig tracks for each tissue, sex, and species; updated annotations for the nine Drosophila species (including *Dmel*, which includes novel isoforms but not novel genes due to it being used as the training dataset) in both gff3 and gtf format. PacBio Iso-seq cDNAs are provided in the SRA (SRP135764). We also provided the updated 1:1 ortholog table between *Dmel* and nonmelanogaster species. Junctions in bam format from HiSAT2 alignments of each sample (by SAMTools) are found at Zenodo (https://zenodo.org/).

## Author contribution

BO and HY designed the experiment. MP, KK, TM, and KK reared and maintained the flies. BO and HY dissected flies. HY and MJ performed RNA-seq library preparation. HY analysed the data. BO and HY wrote the manuscript. All authors read and approved the final manuscript.

## Acknowledgements

We are grateful to Ryan Dale, Justin Fear, Sharvani Mahadevaraju, Terence Murphy, Morgan Park, Harold Smith, Yijie Wang, and the Oliver lab for help and suggestions. The NIDDK Genomics core and the NIH Intramural Sequencing Center performed sequencing. This work utilized the computational resources of the NIH High-Performance Computing (HPC) Biowulf cluster (http://hpc.nih.gov). This research was supported in part by the Intramural Research Program of the NIH, the National Institute of Diabetes and Digestive and Kidney Diseases (NIDDK).

